# Population structure of chum salmon and selection on the markers collected for stock identification

**DOI:** 10.1101/828780

**Authors:** Shuichi Kitada, Hirohisa Kishino

## Abstract

Genetic stock identification (GSI) is a major management tool of Pacific salmon (*Oncorhynchus* Spp.) that has provided rich genetic baseline data of allozymes, microsatellites, and single nucleotide polymorphisms (SNPs) across the Pacific Rim. Here, we analyzed published data sets for adult chum salmon (*Oncorhynchus keta*), namely 10 microsatellites, 53 SNPs, and a mitochondrial DNA locus (mtDNA3, control region and NADH-3 combined) from 495 locations in the same distribution range (*n* = 61,813). TreeMix analysis of the microsatellite loci identified the highest level of genetic drift towards Japanese/Korean populations and suggested two admixture events from Japan/Korea to Russia and the Alaskan Peninsula. The SNPs had been purposively collected from rapidly evolving genes to increase the power of GSI. The highest expected heterozygosity was observed in Japanese/Korean populations for microsatellites, whereas it was highest in Western Alaskan populations for SNPs, reflecting the SNP discovery process. By regressing the SNP population structures on those of the microsatellites, we estimated the selection on the SNP loci according to deviations from the predicted structures. Specifically, we matched the sampling locations of the SNPs with those of the microsatellites according to geographical information and performed regression analyses of SNP allele frequencies on the two coordinates of multi-dimensional scaling (MDS) of matched locations obtained from microsatellite pairwise *F*_ST_ values. The MDS first axis indicated a latitudinal cline in American and Russian populations, whereas the second axis found a differentiation of Japanese/Korean populations. The top five outlier SNPs were mtDNA3 (combined locus of the control region and NADH-3), U502241 (unknown), GnRH373, ras1362, and TCP178, which were consistently identified by principal component analysis. We summarized the functions of the 53 nuclear SNPs and mtDNA3 locus by referring to a gene database system and discussed the functions of the outlier SNPs and fitness of chum salmon.

## 1 INTRODUCTION

Chum salmon (*Oncorhynchus keta*) has a wide distribution range from the Arctic to California in North America, and from Siberia to Japan and Korea in Asia (Salo, 1991). The ability to adapt to the varied environments of spawning rivers in the past and at present may have expanded its distribution range. Chum salmon also comprise the world’s largest hatchery release program (Amoroso et al., 2017; Flagg, 2015; Kitada, 2018; Naish et al., 2007), and every year, more than 3 billion hatchery-reared chum salmon are released in the North Pacific. Therefore, chum salmon is one of the best species to study the structure and history of populations, which could be underpinned by environmental adaptation and anthropogenic selection pressures.

The genetic population structure of chum salmon has traditionally been studied in the context of genetic stock identification (GSI). GSI has been a major management tool of Pacific salmon (*Oncorhynchus* Spp.) since the early 1990s (Beacham et al. 2008a-d, 2009b, 2020; Larson et al., 2014; Shaklee, 1999; Utter, 1991), and it has significantly contributed to high-seas and coastal migration studies (Myers et al., 2007; Seeb et al., 2004). In Pacific salmon GSI, samples from mixed-stock fishery and forensic studies were analyzed with the general objective of providing the optimal resolution of mixing proportions of stocks at a reasonable cost (Beacham et al., 2020). GSI studies have provided genetic baseline data for salmon populations across the Pacific Rim, and these data have contributed to studies into population structure, mixed-stock fisheries, and genetic interactions between hatchery and wild salmon (Waples et al. 2020).

Genetic markers for GSI have progressed from allozymes to microsatellites and single-nucleotide polymorphisms (SNPs) (Beacham et al. 2020; Bernatchez et al., 2017). Allozyme loci often have a small number of alleles. To improve the power of GSI resolution for the high gene flow salmonids, microsatellites were developed because the number of alleles is generally much larger than that of isozymes and much more information can be included. However, standardizing hundreds of microsatellite alleles across sampling points in different countries is difficult (Seeb et al., 2011). To avoid the standardization problem, genotyping of microsatellites of salmon species was generally performed by a single laboratory (Beacham et al. 2008a,b, 2009a,b; Seeb et al., 2011). In contrast, calibrating SNP genotyping is more straightforward because genotype data can be stored in a unified format and can be accessed by different laboratories on different continents (Waples et al. 2020).

Chum salmon is one of the major salmonid species that have been studied for GSI, and SNPs with large allele frequency differences among populations have been genotyped in three different studies to improve the power of GSI (Elfstrom et al., 2007; Smith et al., 2005a,b). The genes that were selected for SNP typing were originally identified as rapidly evolving genes (Elfstrom et al., 2007; Seeb et al., 2011) that showed positive selection in humans and chimpanzees (Nielsen et al., 2005). They included genes that were associated with fatty acid synthesis, testis-specific expression, olfactory receptors, immune responses, and cell growth and differentiation (Elfstrom et al., 2007; Smith et al., 2005a,b). The population structure determined using the SNPs selected for the GSI was biased not only by the selection of the genes but also by the SNP discovery process. Specifically, the three studies were focused on Western Alaska, which was the area of the authors’ interest (Seeb et al., 2011). Consistently, the SNP allelic richness and heterozygosity is high in Alaskan populations.

Neutral and adaptive markers and combinations of them can be useful in establishing optimal management strategies (Funk et al., 2012). Population structures inferred using neutral markers reflect gene flow and genetic drift (Waples & Gaggiotti, 2006), which affect within and among population variations and can lead to adaptive divergence in the genome (Funk et al., 2012). To integrate adaptive markers into the definition of conservation units, Funk and colleagues proposed a framework of comparing population structures inferred from putatively neutral and adaptive loci. The inclusion of information on putatively adapted loci can help to understand mechanisms of local adaptation and is useful for conservation and management of the species (Moore et al., 2014).

Here, we analyzed the published data sets of microsatellites and the SNPs genotyped for chum salmon GSI. First, we inferred the chum salmon population structure and its demographic history using the microsatellite data in a distribution range. Then, we matched the sampling locations of the SNP genotyping studies with those of the microsatellite data according to the available geographical information. By regressing the SNP population structure on the microsatellite population structure, we estimated the selection on the SNPs as deviations from the predicted structure.

## 2 MATERIAL AND METHODS

### 2.1 Screening of population genetics data for chum salmon

We screened population genetics studies of chum salmon in the literature published after 1990 using the Google Scholar search system with keyword searches of “mixed-stock fisheries”, “population structure”, “salmon”, and “stock identification”. We also added studies that were known to us. The population genetics of chum salmon in the North Pacific has been studied extensively over the last four decades. International sampling of chum salmon has been conducted cooperatively across its distribution range to establish baseline genotype data for effective GSI (Beacham et al., 2008a,b, 2009a,b; Seeb & Crane, 1999; Seeb et al., 1995; Urawa et al., 2005, 2009; Wilmot et al., 1994). The population structure of chum salmon has also been studied extensively using various markers, such as isozymes (Kijima & Fujio, 1979; Okazaki, 1982; Sato & Urawa, 2015; Seeb & Crane, 1999; Seeb et al., 1995; Wilmot et al., 1994; Winans et al., 1994), mitochondrial DNA (mtDNA) (Garvin et al., 2010; Park et al., 1993; Sato et al., 2004; Yoon et al., 2008), minisatellites (Taylor et al., 1994), and microsatellites (Beacham et al., 2008a,b, 2009a,b; Olsen et al., 2008. More recently, SNPs were developed for accurate stock identification and have also been used for population genetics (Garvin et al., 2013; Petrou et al., 2014; Saito et al., 2020; Sato et al., 2014; Seeb et al., 2011; Small et al., 2015). A high-throughput panel was developed for chum salmon and patterns of linkage disequilibrium have been examined (McKinney et al., 2020).

Through the data screening, we found several publicly available data sets that covered the distribution range, including Japan, of chum salmon. These data sets comprised microsatellite allele frequencies (Beacham et al., 2009a), nuclear SNP genotypes with a combined mtDNA locus (Seeb et al., 2011), isozyme allele frequencies (Winans et al., 1994; Seeb et al., 1995), and mtDNA control regions (Sato et al., 2004). In this study, we analyzed the microsatellite (Beacham et al., 2009a) and SNP (Seeb et al., 2011) data sets. All Japanese samples were caught in hatcheries and weirs in hatchery-enhanced rivers and were therefore hatchery-reared fish and/or hatchery descendants.

### 2.2 Microsatellite and SNP data sets analyzed in this study

Microsatellite allele frequencies of chum salmon were retrieved from the Molecular Genetics Lab Online Data, Fisheries and Oceans Canada website (https://www.pac.dfo-mpo.gc.ca/science/facilities-installations/pbs-sbp/mgl-lgm/data-donnees/index-eng.html, accessed 01 August 2020). The data consisted of the allele frequencies at 14 loci from chum salmon populations at 381 localities in a distribution range (*n* = 51,355) (Table S1; Figure S1) (Beacham et al., 2009a). These data were used to infer the population structure and demographic history of chum salmon.

SNP genotypes were retrieved from the Dryad data repository (Seeb et al, 2011). The data consisted of 58 SNPs collected from 114 locations in a distribution range (Table S2; Figure S2) (Seeb et al, 2011). We excluded four loci, *Oke_U401-220, Oke_GHII-2943, Oke_IL8r-272*, and *Oke_U507-87*, to avoid pseudo-replication following the original study. A total of 53 nuclear SNPs and a combined mtDNA locus (mtDNA3) were included in our analysis (*n* = 10,458). The mtDNA3 locus contained three gene loci, *Oke-Cr30* and *Oke-Cr386* (mtDNA control region), and *Oke-ND3-69* (mtDNA NADH-3).

### 2.3 Accounting for ascertainment bias in the data sets

Significant deviations from Hardy–Weinberg equilibrium at four loci (*Oke*3, *Ots*103, *One*111 and *OtsG68*) from a 14-microsatellite locus panel were found in Japanese, Russian, and representative North American populations (Beacham et al., 2008a,b). These deviations were likely because of ascertainment bias (Seeb et al., 2011), and therefore we excluded these four loci and used the remaining 10 loci in our analysis.

For the SNP data, the highest levels of allelic richness detected in parts of the Alaskan region may be affected by ascertainment bias (Seeb et al., 2011). Allele/site frequency spectra have been used to examine potential ascertainment bias in SNPs (Nielsen et al., 2004). We calculated allele frequency spectra in six geographical areas according to the major lineage used in the original study (Seeb et al., 2011), namely Western Alaska, Yukon/Kuskokwim upper, Alaskan Peninsula, Southeast Alaska (SEA)/British Columbia (BC)/Washington (WA), Russia, and Japan/Korea.

### 2.4 Inferring population structure and demographic history

We used the microsatellite allele frequencies (Beacham et al., 2009a) to infer the population structure of chum salmon. In previous coalescent simulations, we found that the bias-corrected *G*_ST_ moment estimator (Nei & Chesser, 1983) performed better than other *F*_ST_ estimators when estimating pairwise *F*_ST_ values (Kitada et al., 2017). Therefore, in this study, we used the Nei & Chesser *G*_ST_ moment estimator (NC83) to compute pairwise *F*_ST_. Because only allele frequency data could be accessed, we used the ‘read.frequency’ function and computed pairwise *F*_ST_ values averaged over loci using the ‘GstNC’ function in the R package FinePop ver.1.5.1 on CRAN. A multi-dimensional scaling (MDS) analysis was performed on the pairwise *F*_ST_ distance matrix using the ‘cmdscale’ function in R. We used the cumulative contribution ratio up to the *k*th axis *j*= 1, …, *k*, …, *K*) as the explained variation measure, which was computed in the R function as 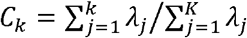, where *λ*_*j*_ is the eigenvalue and *λ*_*j*_ = 0 if *λ*_*j*_ < 0. Following the original study (Beacham et al., 2009a), the populations were classified into eight geographical areas: Japan, Korea, Russia, Alaskan Peninsula, Western Alaska, Yukon/Tanana/Upper Alaska, SEA/BC, and WA (Table S1). A neighbor-joining (NJ) tree (Saitou & Nei, 1987) was constructed using the pairwise *F*_ST_ distance matrix using the ‘nj’ function in the R package ‘ape’.

To infer demographic history with admixture, we applied TreeMix (Pickrell &Pritchard, 2012) for the microsatellite allele frequencies (Beacham et al., 2009a). The analyses were performed using six regional genetic groups, where Japan/Korea and SEA/BC/WA were combined to reduce the number of parameters to be estimated under the limited number of loci. On the basis of the allele frequencies and numbers of repeats, we computed the mean and variance in length at each microsatellite locus and used them to run TreeMix v1.13. We tested up to five migration events and found the best tree was based on the Akaike Information Criterion (AIC; Akaike, 1973); 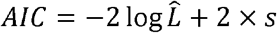, where 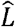, is the maximum composite likelihood and *s* is the number of parameters to be estimated, which, in this study, was three per migration event, namely, migration edge, branch length, and migration weight. For example, for the model with two migration events, the number of parameters was six. TreeMix maximizes composite likelihood and the 2×log likelihood ratio is expected to follow an approximate *χ*^2^ distribution with 3 degrees of freedom. Because TreeMix calculates composite likelihoods, the penalty in AIC needs to account for the correlation in the likelihoods. To avoid unconscious bias towards selection of parameter-rich models, we kept all the candidate models except those where the AIC values were decisively different.

### 2.5 Gene flow and genetic diversity

We recorded approximate longitudes and latitudes of the sampling sites according to the names of rivers and/or areas and maps from the original studies using Google Maps. Sampling locations were plotted on the map using the ‘sf’ package in R. Sampling points with pairwise *F*_ST_ values lower than a given threshold were connected by yellow lines to visualize the population connectivity. Under the assumption of Wright’s island model 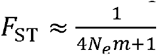, where *N*_*e*_ is the effective population size and *m* is the average migration rate between populations (Slatkin, 1987), a threshold of *F*_ST_ = 0.01 refers to 4*N*_*e*_ *m ≈* 99 migrants per generation (see, Whitlock & Mccauley, 1999; Waples & Gaggiotti, 2006).

We computed expected heterozygosity (*He*) values for the data sets. For microsatellites, we computed *He* for each population at each locus based on the allele frequencies. We then averaged *He* values over loci. For the 53 nuclear SNPs, we converted the original genotype data to genepop format (Rousset, 2008), and read the data using the ‘read.GENEPOP’ function in the R package FinePop2 ver.0.4 on CRAN, where *He* values were computed automatically. The mtDNA3 locus had five alleles, and *He* was computed using the allele frequencies. We also computed NC83 pairwise *F*_ST_ values using 53 nuclear SNPs using the ‘pop_pairwiseFST’ function in FinePop2.

Sampling points were visualized by a color gradient of population-specific *He* values. The color of population *i* was rendered as RGB (1− *H*_*e*0,*i*_, 0, *H*_*e*0,*i*_), where *H*_*e*0,*i*_ = *H*_*e,i*_ − min*H*_*e*_)/ max*H*_*e*_ - min*H*_*e*_). This conversion represents the standardized magnitude of an *He* value at the sampling point, with a continuous color gradient ranging from blue to red for the lowest and highest *He* values, respectively. Population-specific *He* values of microsatellites and SNPs were grouped by seven geographical areas: Japan/Korea, Russia, Alaskan Peninsula, Western Alaska, Yukon/Tanana/Kuskokwim, SEA/BC, and WA. Differences in average *He* values between all pairs of the groups were tested by one-way ANOVA, where *p*-values were adjusted for the multiple comparisons using the ‘TukeyHSD’ function in R.

### 2.6 Identifying highly differentiated SNPs beyond the neutral population structure

To compare the genetic differentiation of SNP allele frequencies with the population structure inferred using the microsatellite markers, we analyzed 10 microsatellite loci and 53 nuclear SNPs loci plus the combined mtDNA3 locus. The mtDNA3 locus had five alleles and the major allele (second one) was used for the meta-analysis. The functions of all 54 analyzed gene loci were confirmed by referring to the GeneCards database system (https://www.genecards.org/) and published literature.

First, we matched the 114 sampling locations of the SNPs (Figure S2) to the nearest location of the 381 sampling points of the microsatellites (Figure S1). The same locations were not necessarily used for genotyping. When we did not find the same location names, we assigned the closest locations of the microsatellites to those of SNPs using the longitudes and latitudes of the sampling locations. Among the 114 sampling locations for SNPs, 56 locations also had samples for microsatellite genotyping. We assigned the closest locations of microsatellites to the other 58 locations for SNPs. Then, we identified loci with allele frequencies that were significantly differentiated from the microsatellite population structure in the pairwise F_ST_. Specifically, we performed regression analyses of allele frequencies of the 53 nuclear SNPs and the mtDNA3 locus on the two coordinates (axes) of the MDS of the pairwise F_ST_ estimated using the microsatellite allele frequencies as:

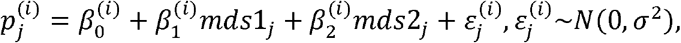

where, 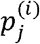 is the allele frequency of SNP *i*(= 1, …, *l*) in population *j* (= 1, …, *J*). The explanatory variables *mds*1_*j*_ and *mds*2_*j*_ are the 1^st^ and 2^nd^ MDS coordinates of pairwise *F*_ST_ values estimated from the microsatellite allele frequencies for the matched populations *j* (= 1, …, *J*). These explanatory variables are population specific (*J* = 114). The regression analysis was performed for each marker (*i*= 1, …, 54) and 54 sets of regression coefficients 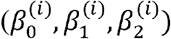 were obtained. The *p*-values for the coefficients were adjusted for the multiple comparison (*q*-values) by the method of Benjamini & Hochberg (1995) using the ‘p.adjust’ function in R. Given the population structure predicted from the neutral microsatellites, the outlier loci from the set of 53 nuclear SNPs and mtDNA3 locus that deviated largely from the predicted population structure are likely to be under natural selection and/or anthropogenic selection.

To identify genes that best characterize geographical areas, we conducted a principal components analysis (PCA) for the allele frequencies of the 53 nuclear SNPs and mtDNA3 locus using the ‘prcomp’ function in R. Finally, we visualized the geographical distributions of the identified highly differentiated genes using boxplots and geographical maps of the allele frequencies and *He* values. The sampling locations were mapped with a color gradient of allele frequencies for population *j* rendered as RGB (*p*_*j*0_, 0, 1 − *p*_*j*0_), where *p*_*j*0_ = (*p*_*j*_ − min *p*)/ max *p* − min *p*). This conversion represents the standardized allele frequency values at the sampling point, with a continuous color gradient ranging from blue to red for the lowest and highest allele frequencies, respectively.

## 3 RESULTS

### 3.1 Population structure and demographic history of chum salmon

Pairwise *F*_ST_ values based on the microsatellite allele frequencies were 0.019 ± 0.010 (SD, standard deviation). The MDS of pairwise *F*_ST_ values (Figure 1a) indicated a latitudinal cline in the American and Russian populations and a separation of Alaskan populations from the others along the first axis (mds1) explaining 29% of the variance. The second axis (mds2, 16% of variance) showed a differentiation of Japanese/Korean populations from the others, although they were closely related to southern Russian populations from Sakhalin, Amur, and Primorye. Plots of mds1 vs mds3, and mds1 vs mds4 produced similar results, showing most differentiation was driven by divergence from a latitudinal cline (Figure S3). The mds1 to mds4 explained 60% variation in the pairwise *F*_ST_ distant matrix (Figure S4). In the unrooted NJ tree generated from this data set (Figure S5a), five large regional population groupings were inferred: (i) Alaskan Peninsula, (ii) Western Alaska/Yukon (Canada), (iii) Russia/Japan/Korea, (iv) SEA/Northern BC, and (v) Southern BC/WA. The TreeMix analysis indicated two periods of admixture (migration events), from Japan/Korea to Russia and the Alaskan Peninsula (Figure 1b; Figure S6). Japanese/Korean populations had the highest level of genetic drift among all the groups, whereas the drift was the lowest in the Alaskan Peninsula and SEA/BC/WA populations.

**FIGURE 1.**
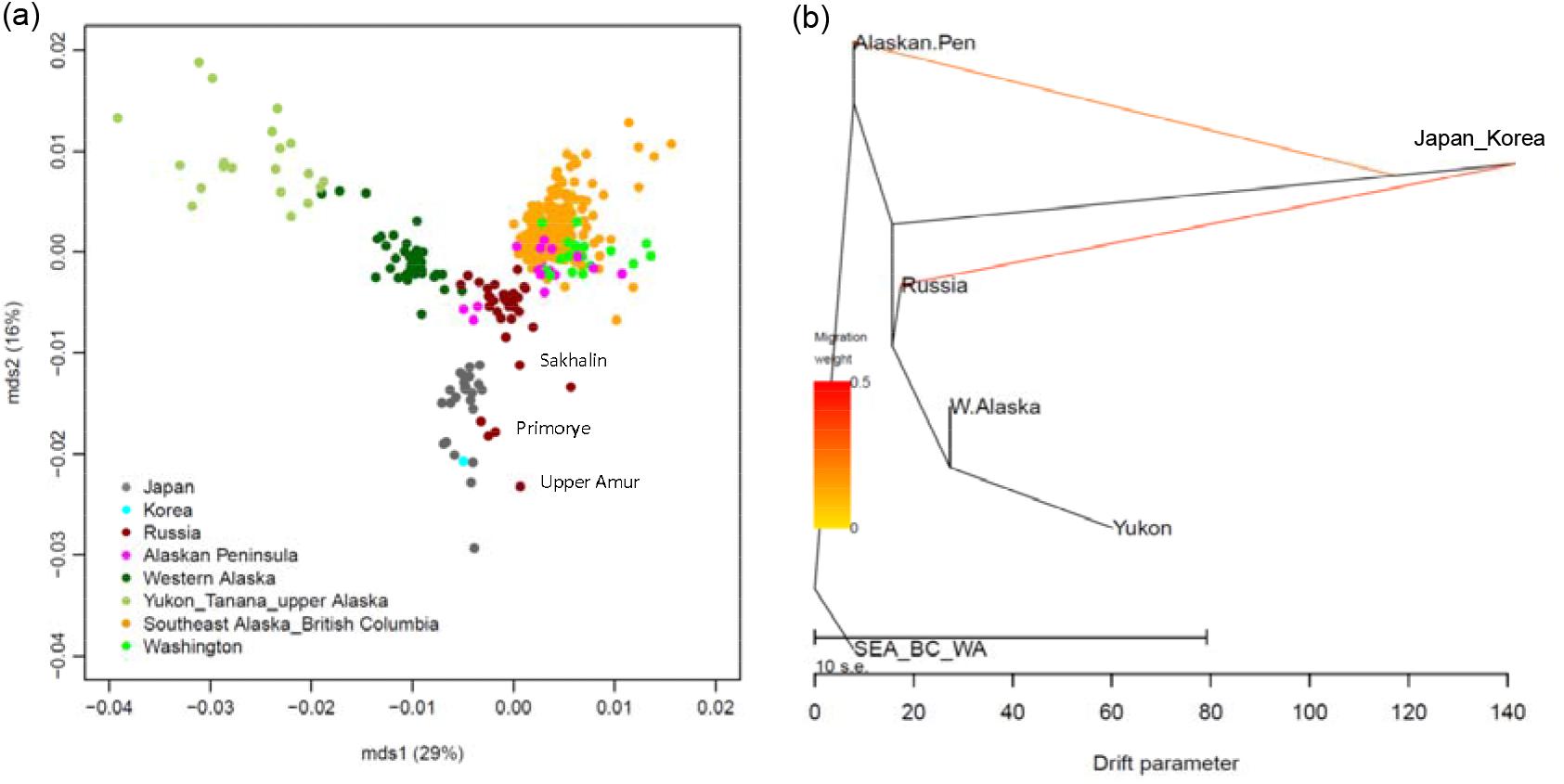
Population structure and admixture events of chum salmon based on microsatellite markers. (a) Multi-dimensional scaling (MDS) plots (mds1 vs mds2) of the population structure of chum salmon based on pairwise *F*_ST_ values estimated from 10 microsatellite loci in 381 populations. (b) Admixture graph inferred using TreeMix with two migration events. The scale bar (drift parameter) is 10 times the average standard error of the sample covariance matrix of allele frequencies between populations (Pickrell & Pritchard, 2012).

### 3.2 Ascertainment bias in the SNP data

The allele frequency spectra (Figure S7) showed a similar pattern in all areas except Western Alaska, where a much lower frequency was found for the lowest allele frequency in Western Alaska.

### 3.3 Gene flow and genetic diversity

Using the criterion of pairwise *F*_*ST*_ <0.01 (4*N*_*e*_ *m* ≈ 99), substantial gene flow between American and Asian populations was identified for the microsatellites (Figure 2a). Japanese populations were found to be connected to Russian and Korean populations. For the SNPs, using the same criterion, we detected substantial gene flow within Alaska, but the SEA and WA populations appeared to be isolated and gene flow was very little between American and Russian populations (Figure 2b). Japanese and Korean populations were isolated from all the others.

**FIGURE 2.**
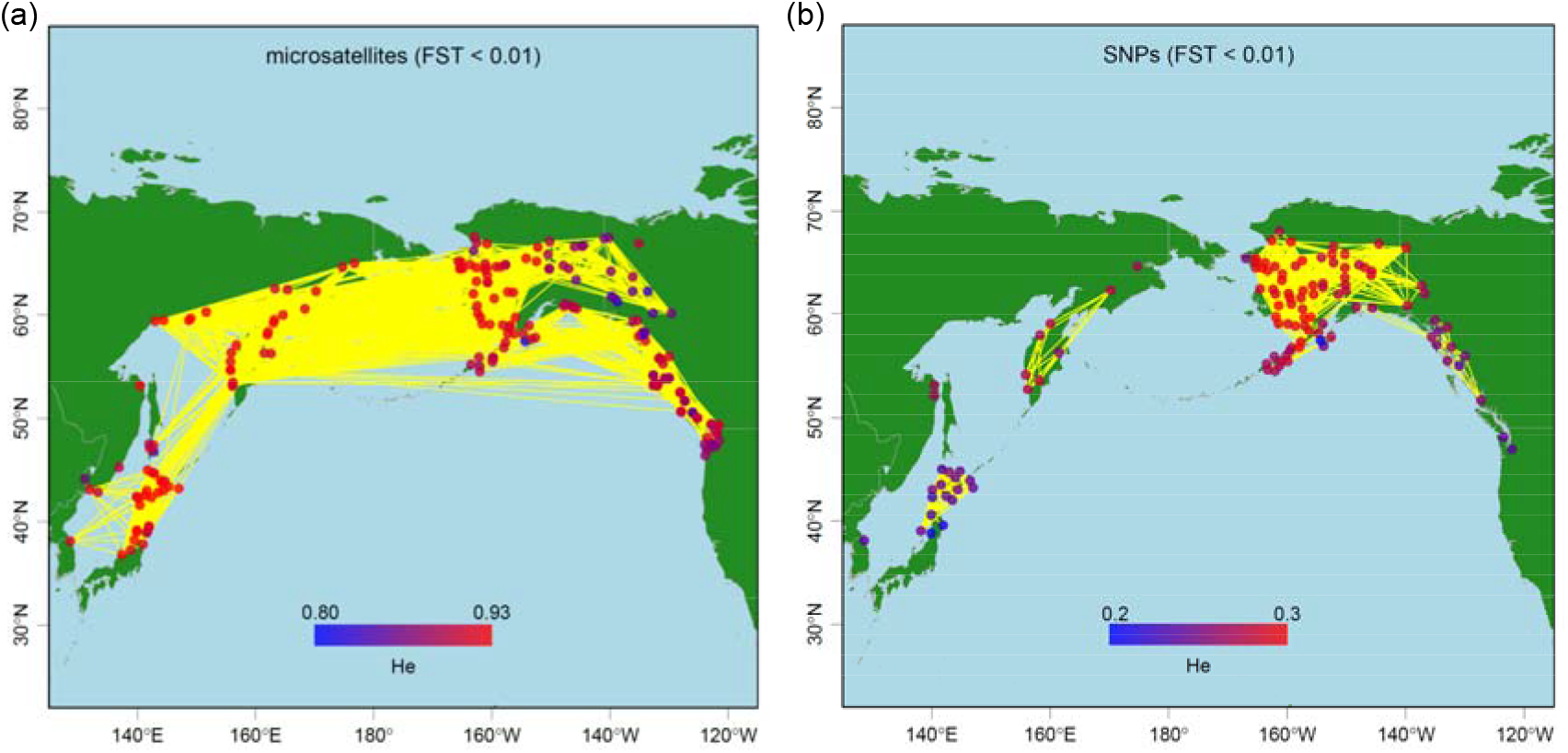
Visualization of genetic diversity and population connectivity of chum salmon based on microsatellite and SNP markers. (a) Map obtained using 10 microsatellite loci of 381 populations (*n* = 51,355) (Beacham et al., 2009a). (b) Map obtained using 53 SNPs of 114 populations (*n* = 10,458) (Seeb et al., 2011). The color of each population reflects the magnitude of the expected heterozygosity (*H*_*e*_) values, with a continuous color gradient ranging from blue to red for the lowest and highest *He* values, respectively. The yellow lines connect populations with pairwise *F*_ST_ <0.01.

For the microsatellites, the *He* values were generally high in almost all the populations (Figure 2a, Figure S8a). The average *He* value over loci was 0.89±0.02 (range 0.81–0.93). High *He* values were found in Japanese, Russian, and Alaskan populations, whereas *He* values were slightly lower in SEA/BC, WA, and Canadian Yukon populations (Figure S5b). Japanese populations had the highest *He* values, but the mean was similar to those in Russian and Western Alaskan populations, lower in Alaskan Peninsular, SEA/BC, and WA populations, and extremely low in the Yukon population (Figure S8a).

In contrast, for the SNPs, the highest *He* values were found in Western Alaska populations, followed by Yukon, Alaskan Peninsula, Russia, and SEA populations (Figure 2b, Figure S8b). The average *He* value over loci was 0.27±0.02 (range 0.20–0.30). Japan and WA populations had the lowest *He* values. The *He* values were similar in populations from Western Alaska, Yukon, Alaskan Peninsula, and Russia (Figure S8b) and, although Russian populations had higher *He* values than the SEA/BC and WA populations, the differences between these populations were not significant. Japanese populations had the lowest *He* values with no significant difference between it and the WA populations.

### 3.4 Differentiated SNPs beyond the neutral population structure

We summarized 53 nuclear SNPs and the mtDNA3 locus with functions (if known), locus names, and raw data locus names used in our analysis (Table S3). Five of the outlier SNPs were identified as potential candidates for selection in the original study (Seeb et al., 2011). We found 31 SNPs in functional genes and 23 SNPs were unknown.

The results of our regression analysis are summarized in Table 1. Most of the SNPs had large coefficients for both mds1 and mds2 with highly significant *q*-values, showing that most of SNPs were differentiated beyond the neutral population structure. The top five outlier loci (Table 1) with the highest regression coefficients and *q*-values were mtDNA3, U502241, GnRH373, ras1362, and TCP178, which characterized mds2 of the pairwise *F*_ST_. The scatter plot of mds1 and mds2 (Figure 3a) identified the outlier SNPs that diverged beyond the scale of mds1. The PCA (Figure 3b) identified differences between Japanese/Korean and other populations as a primary component (PC1, 43% variance), whereas the second component (PC2, 20% variance) corresponded to the latitudinal cline among the Russian/American populations. Biplots of PCA1 versus PCA3 and PCA4 described outliers (Susitna, and Sturgeon River on Kodiak Island), but most differentiation was caused by divergence of the Japanese/Korean populations (Figure S9). PCA1 to PCA4 explained 76% of variance of the SNP allele frequencies (Figure S10). The PCA (Figure 3b) confirmed that differentiation of Japanese/Korean populations were characterized mainly by the top five outlier SNPs detected by the regression analysis (Figure 3a).

**Table 1.**
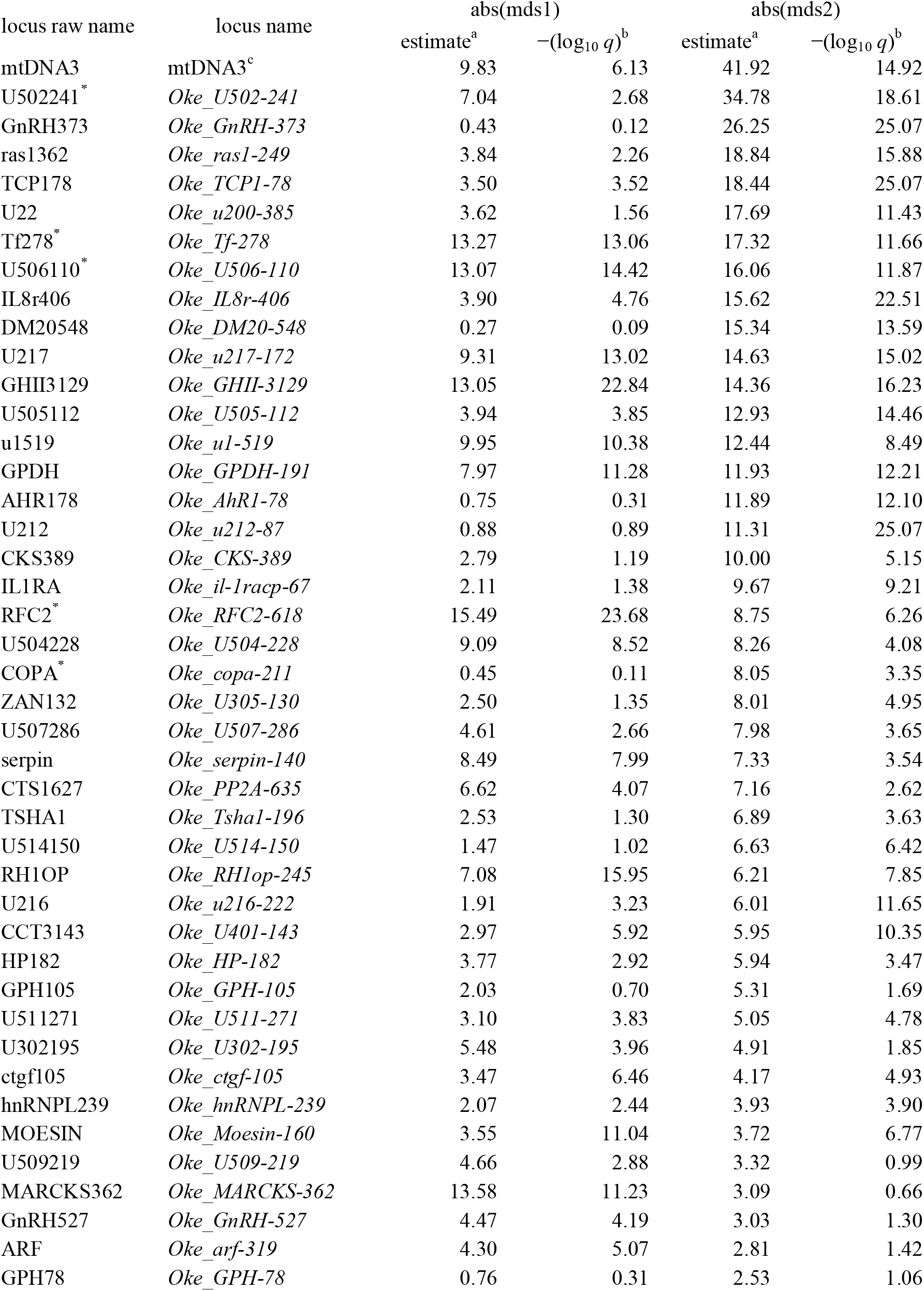
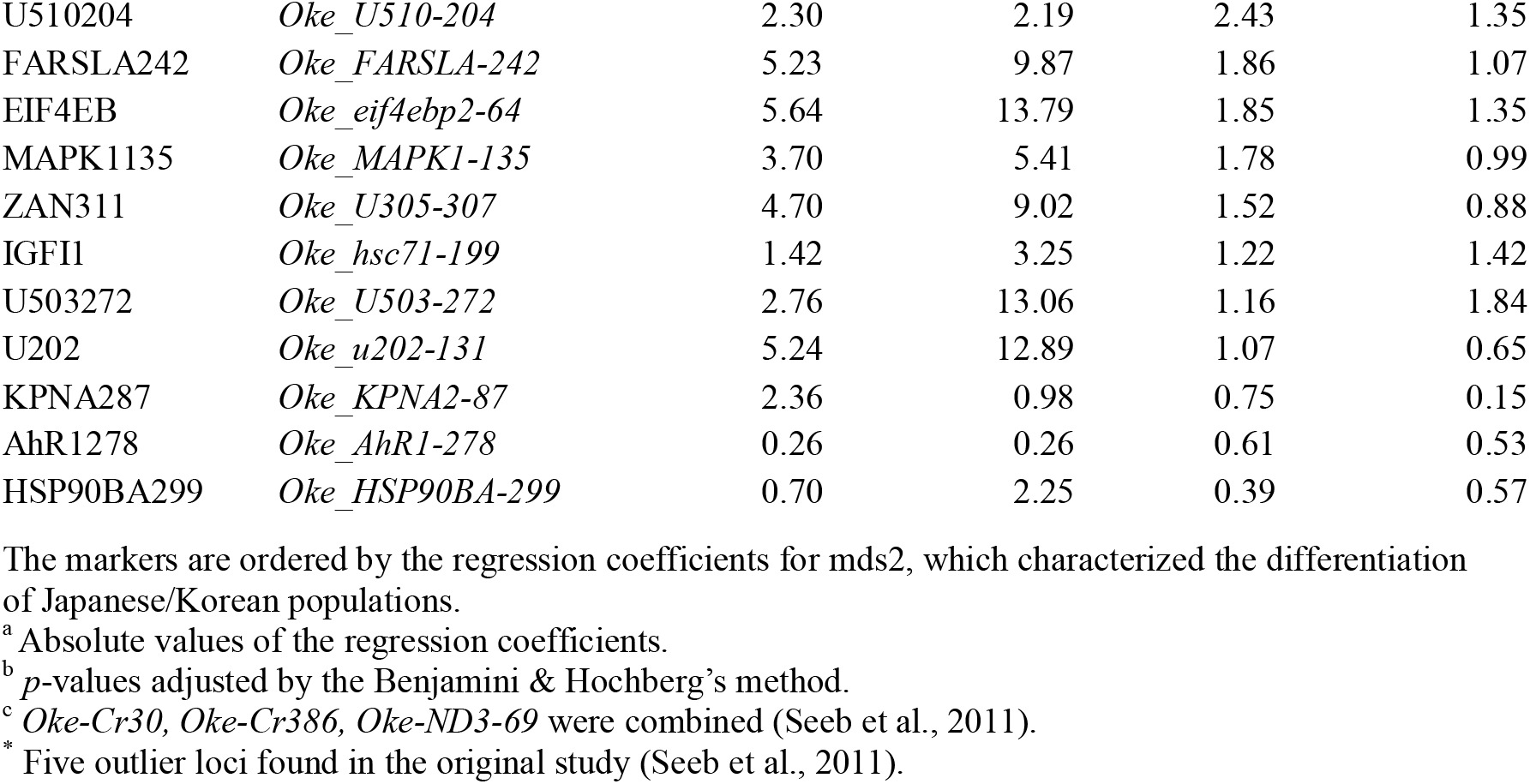
Regression analysis of SNP allele frequencies on abs(mds1) and abs(mds2) of the microsatellite pairwise *F*_ST_ values

**FIGURE 3.**
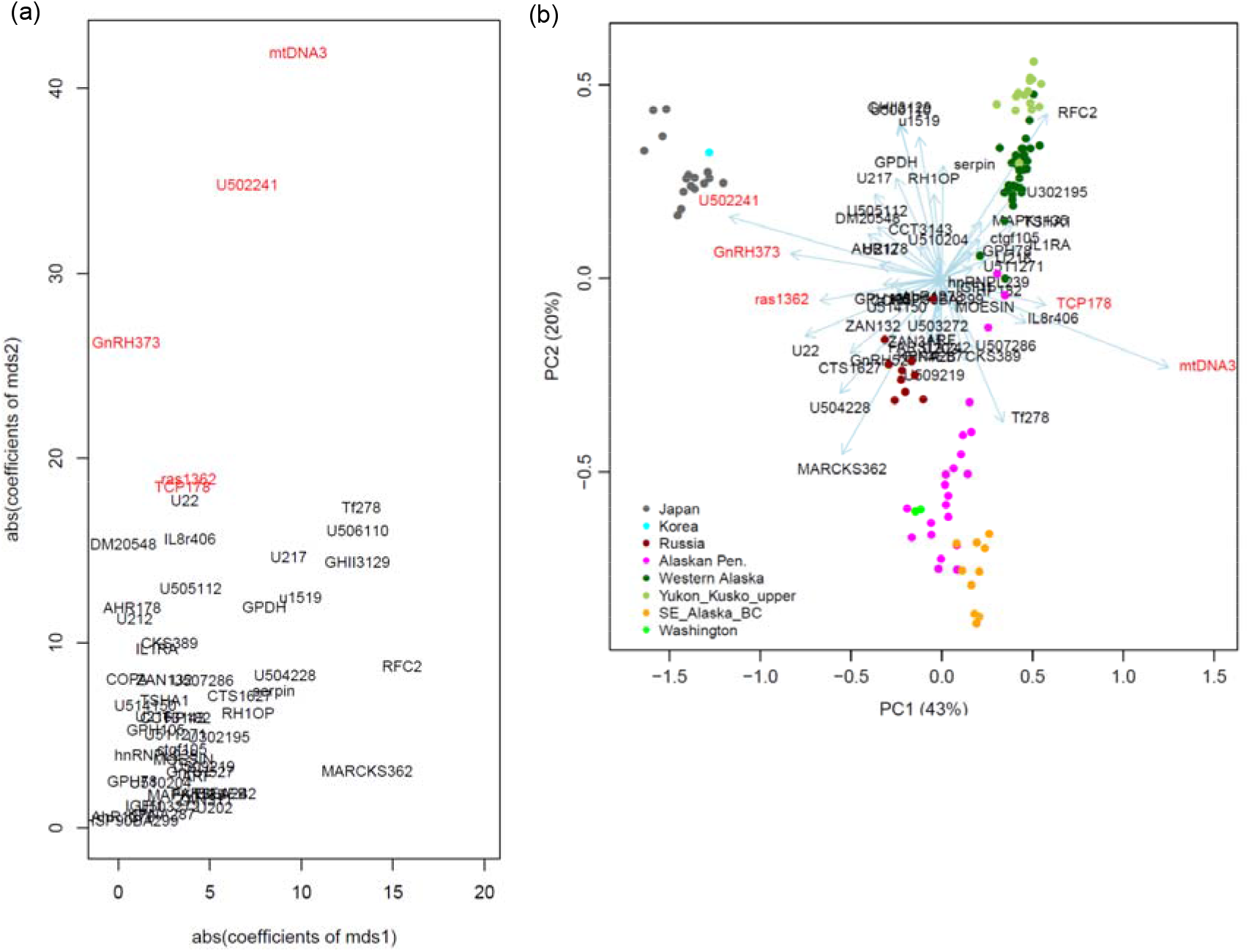
SNPs that characterized the distinctiveness of Japanese/Korean chum salmon populations. (a) Absolute values of the regression coefficients of mds1 vs those of mds2 on the same scale. Red indicates the outlier SNPs that highly diverged beyond the neutral population structure. (b) Principal component analysis (PCA) biplot, based on 53 nuclear SNPs and the combined mtDNA3 locus.

The allele frequencies of the top five outlier loci were distinct in Japanese/Korean populations in the distribution range (Figure 4). The major allele of mtDNA3 was fixed at close to 1.0 in American and Russian populations, whereas the allele frequencies were significantly lower (0.21± 0.11) in Japanese/Korean populations. The U502241 allele frequencies were much higher in Japanese/Korean and Washington populations than they were in the other populations. GnRH373, ras1362, and TCP178 had similar geographical distributions of allele frequencies, and in Japanese/Korean populations they were differentiated from those in the other populations. The average genetic diversity (*He*) was significantly higher in Japanese/Korean populations than that it was in Western Alaska for mtDNA3 (Welch two sample *t* test, *t* = 30.5, df = 17.5, *p* = 2.2 × 10^−l6^) and TCP178 loci (t = 36.8, df = 51.3, *p* = 2.2 × 10^−l6^), whereas it was similar in Western Alaska for U502241 (*t* = 0.9, df = 19.7, *p* = 0.37), and ras1362 (*t* = 0.6, df = 35.7, *p* = 0.57). GnRH373 had lower *He* values in Japanese/Korean populations (0.33) than in Western Alaskan populations (0.40) (*t* = -3.0, df = 21.2, *p* = 0.006).

**FIGURE 4.**
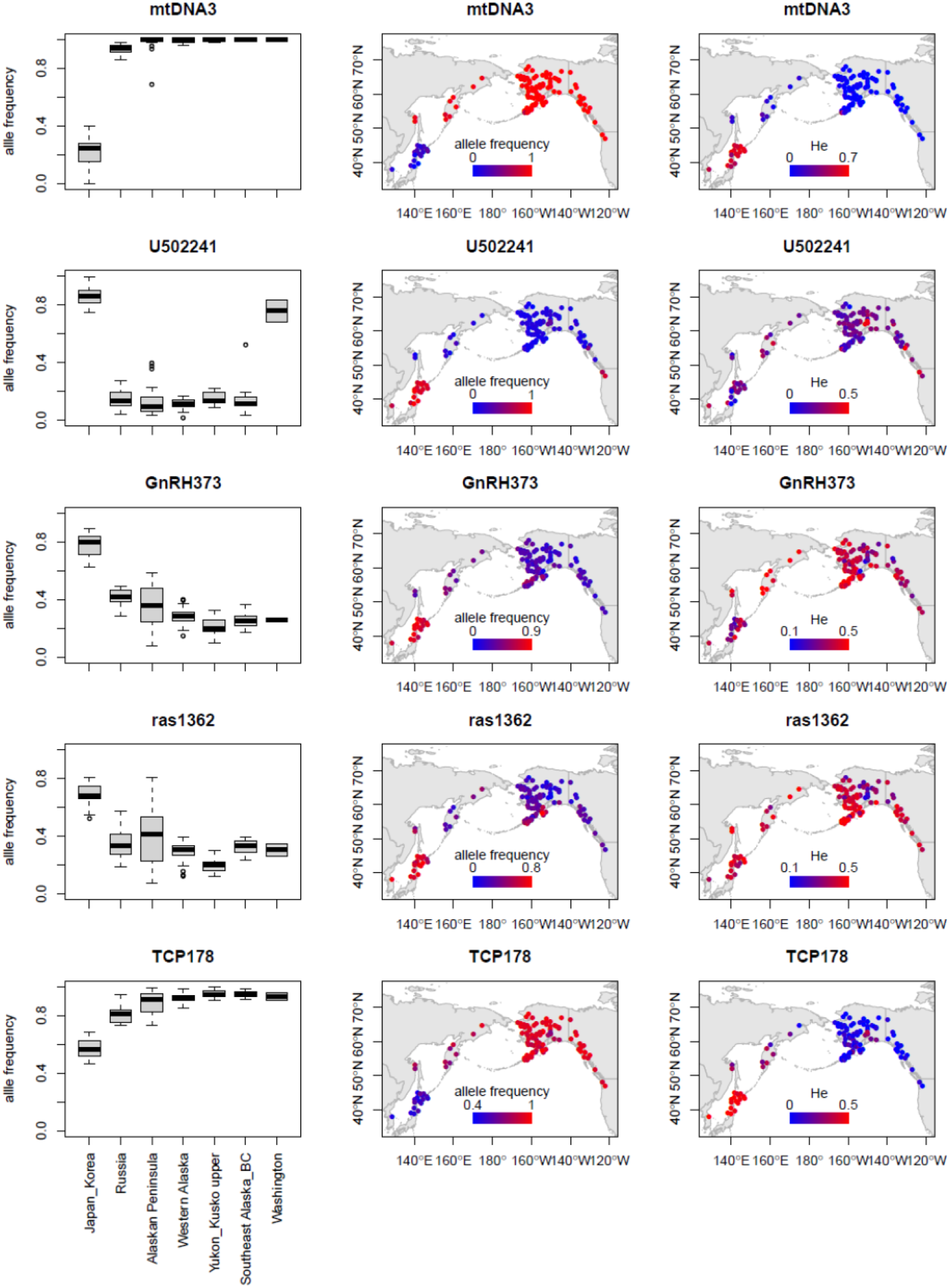
Geographical distribution of the top five outlier SNPs that characterized the distinctiveness of Japanese/Korean chum salmon populations. The points on the maps show allele frequencies (center) and expected heterozygosity (*He*) (right) at sampling locations, with a continuous color gradient ranging from blue to red for the lowest and highest *He* values, respectively.

## 4 DISCUSSION

The inferred population structure based on microsatellites was consistent with the one obtained by the TreeMix analysis (Figure 1). The mds1 of pairwise *F*_ST_ identified the north–south cline in American and Russian populations, and mds2 clearly showed divergence of Japanese/Korean populations, in agreement with the original study (Beacham et al., 2009a). TreeMix indicated that modern chum salmon populations may be recolonized from source populations between the Alaskan Peninsula and SEA/BC/WA, and expanded to the north and to Asia. However, a higher level of drift among the Japanese/Korean populations was clear relative to the Russian and American populations, suggesting that the Japanese/Korean populations were the most diverged from the ancestral population and that they diverged between Russia and the Alaskan Peninsula. Two inferred admixture events may reflect the stray of Japanese/Korean populations into Russian and Alaskan rivers.

The SNP allele frequency spectra (Figure S7) showed a similar pattern in all areas except Western Alaska. Because the entire range was represented for the SNP selection from Japan to WA (Seeb et al., 2011), ascertainment bias caused by cut-off of minor alleles may have occurred in Western Alaska, as indicated in Figure S7. This bias may have increased in intermediate allele frequencies and genetic diversity in the Western Alaska samples relative to other samples, leading to an increase in the genetic diversity of SNPs in the Western Alaska samples (Figure S8b), although the effects of the ascertainment bias were expected to be minimal within Alaska (Seeb et al., 2011).

Our regression analysis identified the top five outlier gene loci (Table 1, Figures 3, 4) that were significantly differentiated beyond the neutral population structure in Japanese/Korean populations, namely, mtDNA3, U502241, GnRH373, ras1362, and TCP178. The scatter plot of mds1 and mds2 indicated that most of the SNPs were differentiated beyond the neutral population structure. Selection can be induced by natural and anthropogenic pressures. The mds1 described the north–south cline of population expansion (Figure 1). The maximum regression coefficient of mds1 was 15.5 for RFC2 (Table 1). In contrast, the top five outlier gene loci had mds2 coefficients >18.4, which suggested that the allele frequencies of the top five gene loci were the result of anthropogenic selection pressures in Japanese/Korean populations.

mtDNA3 (a combined locus of the control region and NADH-3, Smith et al., 2005b) was the most differentiated gene locus (Figures 3, 4). mtDNA encodes some of the proteins in the oxidative phosphorylation enzymatic complex and plays a key role in aerobic ATP production, by contributing to the ability to respond to endurance exercise training (reviewed by Stefàno et al., 2019). The mtDNA control region functions in dNTP metabolism (Nicholls & Minczuk, 2014) and oxygen consumption (Kong et al., 2020). Subunits of the NADH dehydrogenase complex encoded by mtDNA are involved in the pathway from NADH to oxygen (Rivera et al., 1998). A significant reduction in mtDNA3 allele frequencies might reduce the efficiency of aerobic exercise ability and endurance, energy metabolism, and oxygen consumption. These results are consistent with the lower swimming endurance of Japanese hatchery-born chum salmon fry measured in a stamina measurement experimental device in the Japanese oldest Chitose Hatchery, Hokkaido (Kobayashi & Ohkuma, 1983). Wild chum salmon fry had a 1.4-fold higher swimming ability (56.6 ± 11.1 cm/s) than hatchery-reared fry (41.4 ± 12.3) (*t* = 2.45, df = 8.7, *p* = 0.038) in the early 1980s.

U502241 allele frequencies were significantly higher in Japanese/Korean and Washington populations (Figure 4). Elevated levels of U502241 allele frequencies in Japanese/Korean and Washington populations were similar (*t* = 1.3, df = 1.1, *p* = 0.39), suggesting a common effect. The original study (Seeb et al. 2011) found this phenomenon and detected U502241 as an outlier SNP. No function has been associated with the chum U502241 locus so far (Elfstrom et al., 2007), but the Atlantic salmon immunoglobulin IgH locus B genomic sequence was the most likely match for U502241 (Seeb et al. 2011). Immunoglobulins initiate immune responses (Gene Cards).

GnRH (gonadotropin-releasing hormone) is a key regulator of vertebrate reproduction, including that of salmonids (Khakoo et al., 1994). TCP1 (T-Complex 1) is related to sperm–zona–pellucida interaction and spermatozoan fertilization ability (Dun et al., 2011). The *ras* gene has a central role in cell growth (Rotchell et al., 2001). These outliers related to reproduction and growth are key parameters in hatchery production. In chum salmon, GnRH is involved in gonadal maturation during the early and final phases of upstream migration (Kudo et al., 1996). GnRH also improved stream odor discrimination in adult fish (Ueda, 2019), and expression of GnRH increased in the brain during homing migration (Ueda et al., 2016). The differentiation of GnRH373 (Figure 4) might influence homing migration timing, potentially leading to the current introgression from Japanese into Russian populations, as suggested by our TreeMix analysis.

Apparently different distributions in *He* values were observed in microsatellites and SNPs in the Japanese/Korean populations (Figure S8). In the Japanese populations, the highest microsatellite *He* values may have been influenced by the transplantation history in Japanese river populations (Beacham et al., 2008a), whereas the lowest SNP *He* values may be a consequence of artificial selection in the hatcheries. The *He* values in Japanese/Korean and Western Alaskan populations were similar for U502241 and ras1362, whereas the *He* values for GnRH373 were smaller in the Japanese/Korean populations than in the Western Alaskan populations as shown in Figure 4. In contrast, the *He* values in Japanese/Korean populations were significantly higher than those in Western Alaskan populations for mtDNA3 and TCP178, suggesting increases in heterozygosity. These results suggested genotype-specific effects on fitness.

The analyses in this study relied on limited data sets. Clearly, further genomic studies are needed to obtain more precise views of the demographic history, and environmental and anthropogenic selection of chum salmon. Population genomics studies can also provide finer resolution of GSI and will be useful for management and conservation of chum salmon.

## ACKNOWLEDGEMENTS

This research was made possible by the SNP genotypes of chum salmon throughout the distribution range maintained by Lisa W. Seeb and her colleagues, and by the North Pacific baseline microsatellite allele frequencies maintained by Terry D. Beacham and his colleagues at Fisheries and Oceans Canada. We also appreciate researchers and staff for their substantial efforts to obtain baseline samples. This study was supported by the Japan Society for the Promotion of Science Grants-in-Aid for Scientific Research KAKENHI (nos. 18K0578116 to SK and 19H04070 to HK).

## DATA AVAILABILITY STATEMENT

The authors affirm that all data necessary for confirming the conclusions of this article are present within the article, figures, and supporting information.

## SUPPORTING INFORMATION

Additional supporting information may be found online in the Supporting Information section.

