## Supplemental Figures for "Population structure of chum salmon and selection on the markers collected for stock identification"

This PDF file includes Supplemental Figures S1–S10.
Supplemental Tables S1-S3 are in an Excel file.

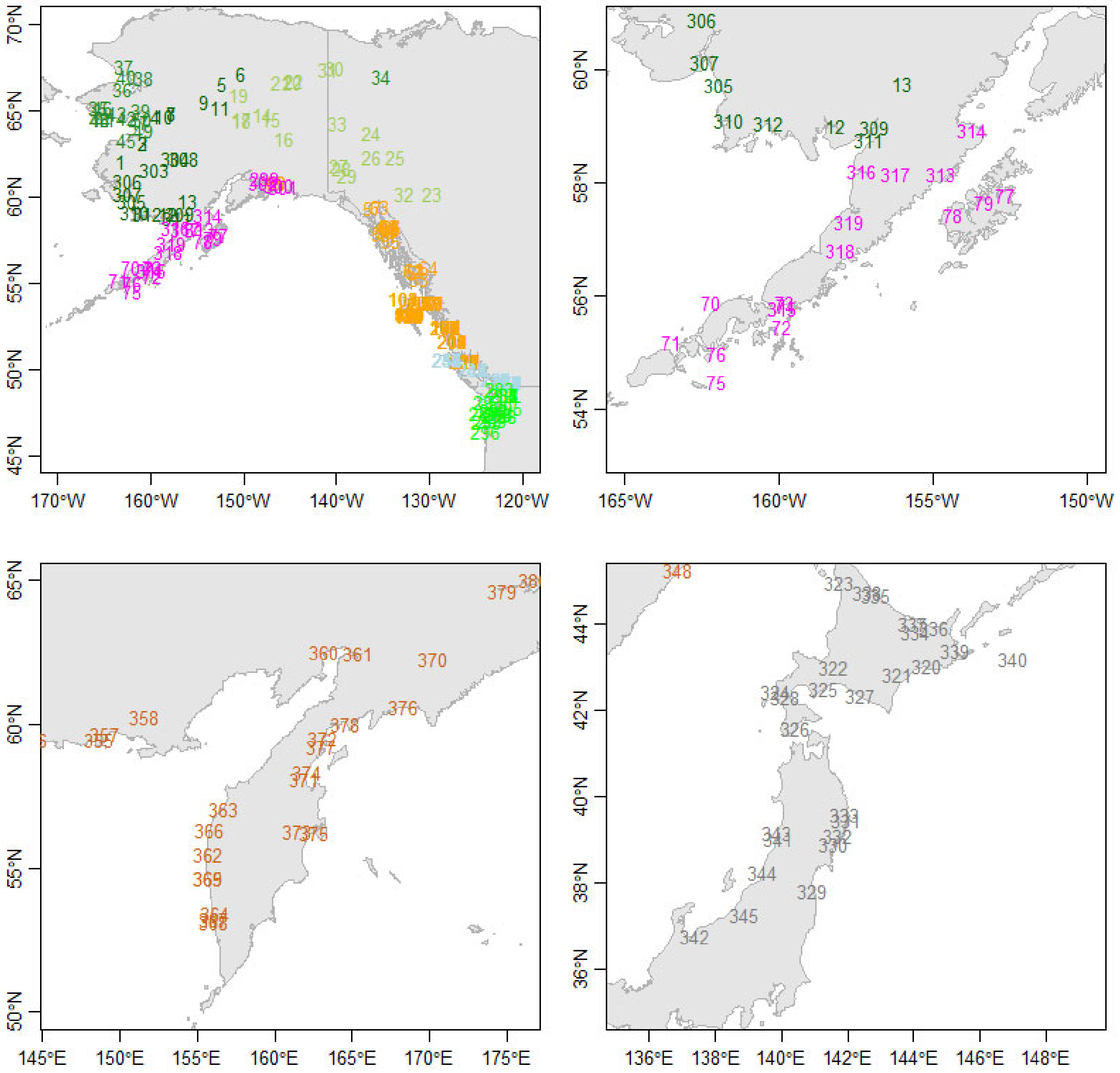


**Figure S1** Sampling locations for chum salmon microsatellite genotyping. Samples were collected in 1986–2007 from 381 populations across the Pacific Rim ($n$ = 51,355). Data are from Beacham et al. (2009a) and are given in Table S1.

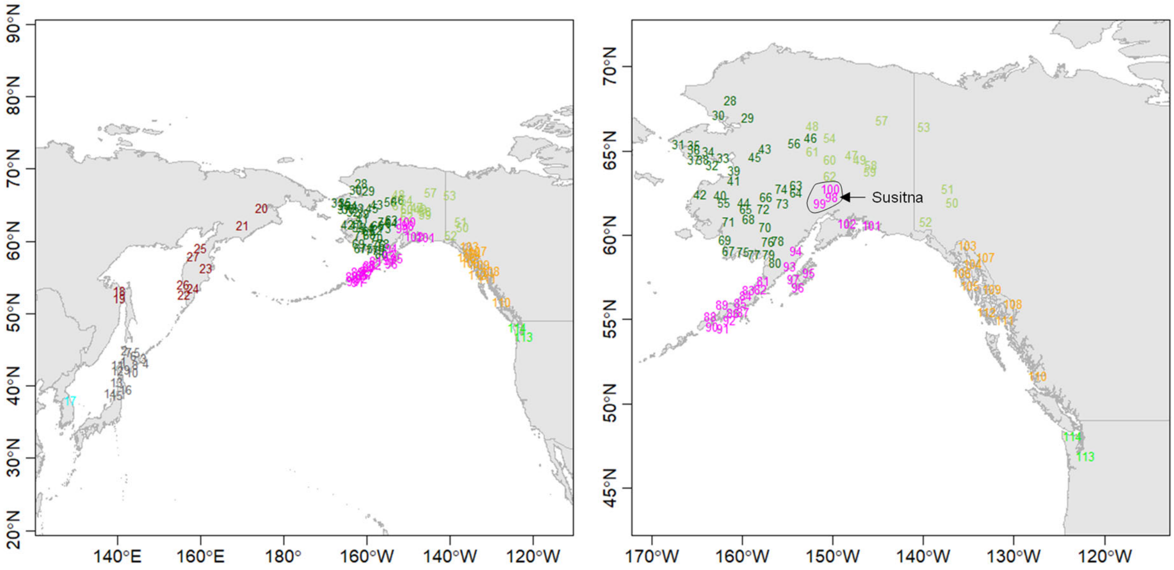


**Figure S2** Sampling locations for chum salmon SNP genotyping. The entire area and North America are shown. Samples were collected in 1989–2006 from 114 populations across the Pacific Rim ($n$ = 10,458). Data are from Seeb et al. (2011) and are given in Table S2.

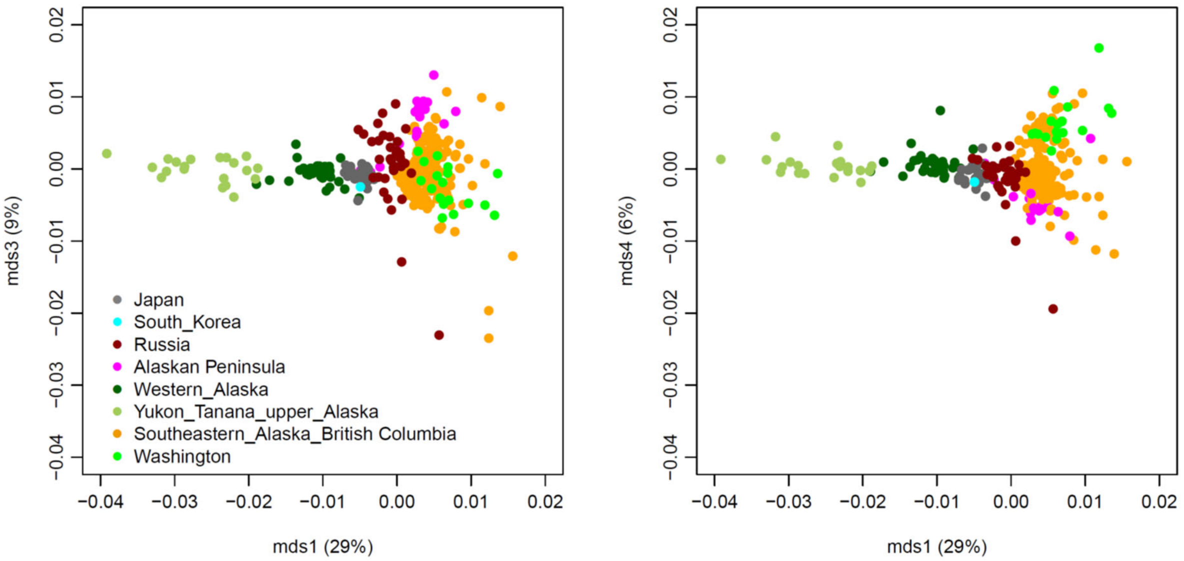


**Figure S3** Multi-dimensional scaling (MDS) plots (mds1 vs mds3, and mds1 vs mds4) of pairwise $F_{ST}$ values of chum salmon in the distribution range estimated from 10 microsatellite loci in 381 populations. Data are from Beacham et al. (2009a) and are given in Table S1.

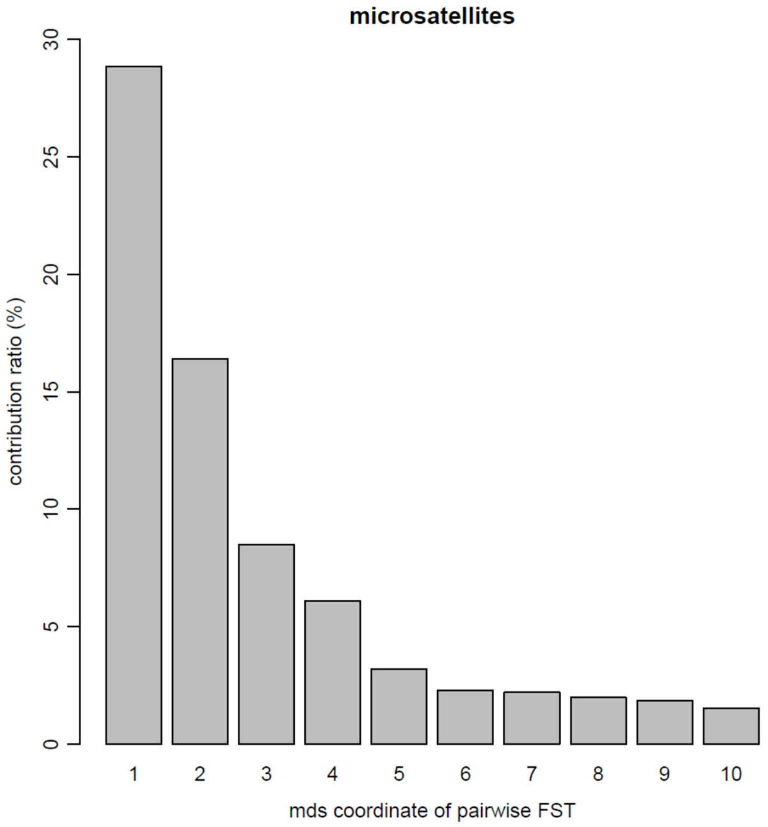


**Figure S4** Contribution of multi-dimensional scaling (MDS) axes of pairwise $F_{ST}$ values estimated from 10 microsatellite loci in 381 chum salmon populations. Data are from Beacham et al. (2009a) and are given in Table S1.

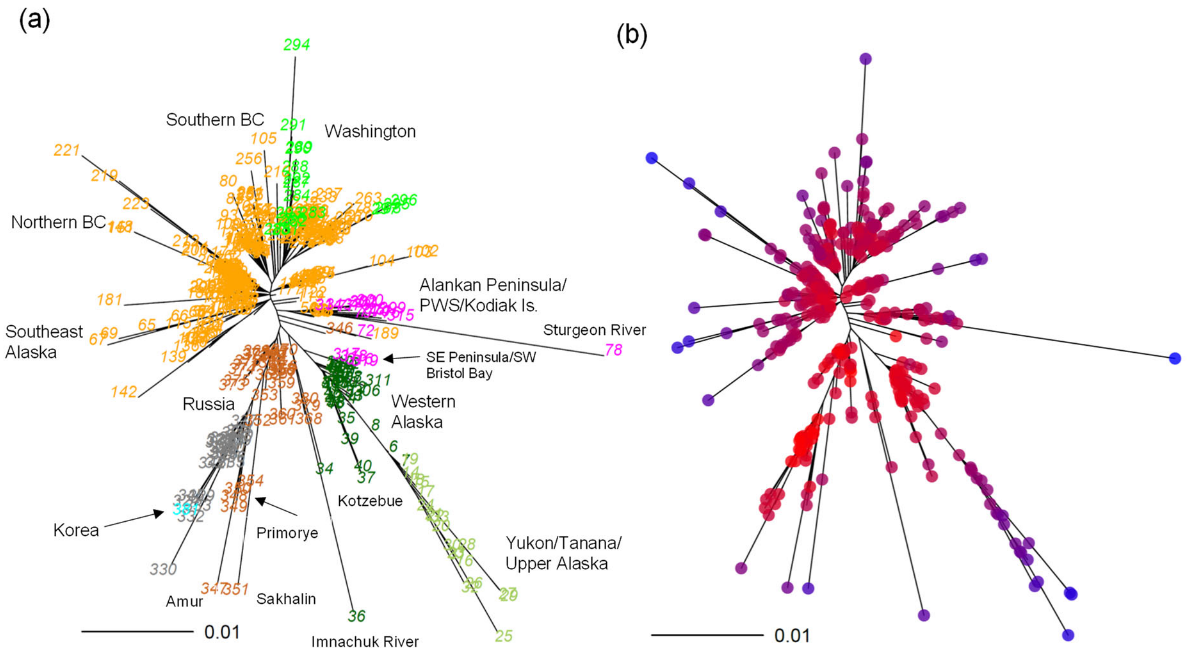


**Figure S5** Population structure and genetic diversity of chum salmon based on 10 microsatellite markers. (a) Unrooted neighbor-joining (NJ) tree based on pairwise $F_{ST}$ values estimated from 381 populations ($n$ = 51,355). Numbers refer to the sampling locations (Table S1, Figure S1). (b) Unrooted NJ tree based on pairwise $F_{ST}$ values overlaid with expected heterozygosity ($H_e$) values. The color of each population reflects the magnitude of the $H_e$ values, with a continuous color gradient ranging from blue to red for the lowest and highest *He* values, respectively. Data are from Beacham et al. (2009a) and are given in Table S1.

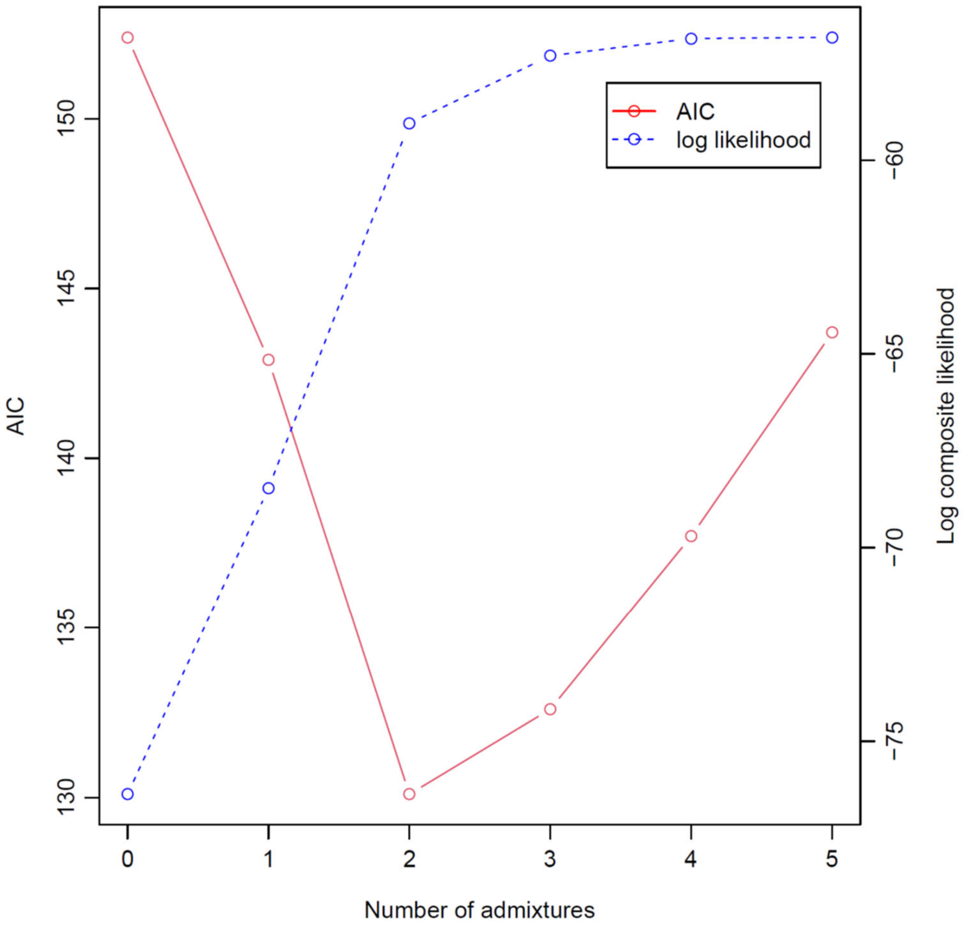


**Figure S6** Composite likelihood and Akaike Information Criterion (AIC) values for models of the number of admixture event from TreeMix based on 10 microsatellite loci in 381 chum salmon populations. Data are from Beacham et al. (2009a) and are given in Table S1.

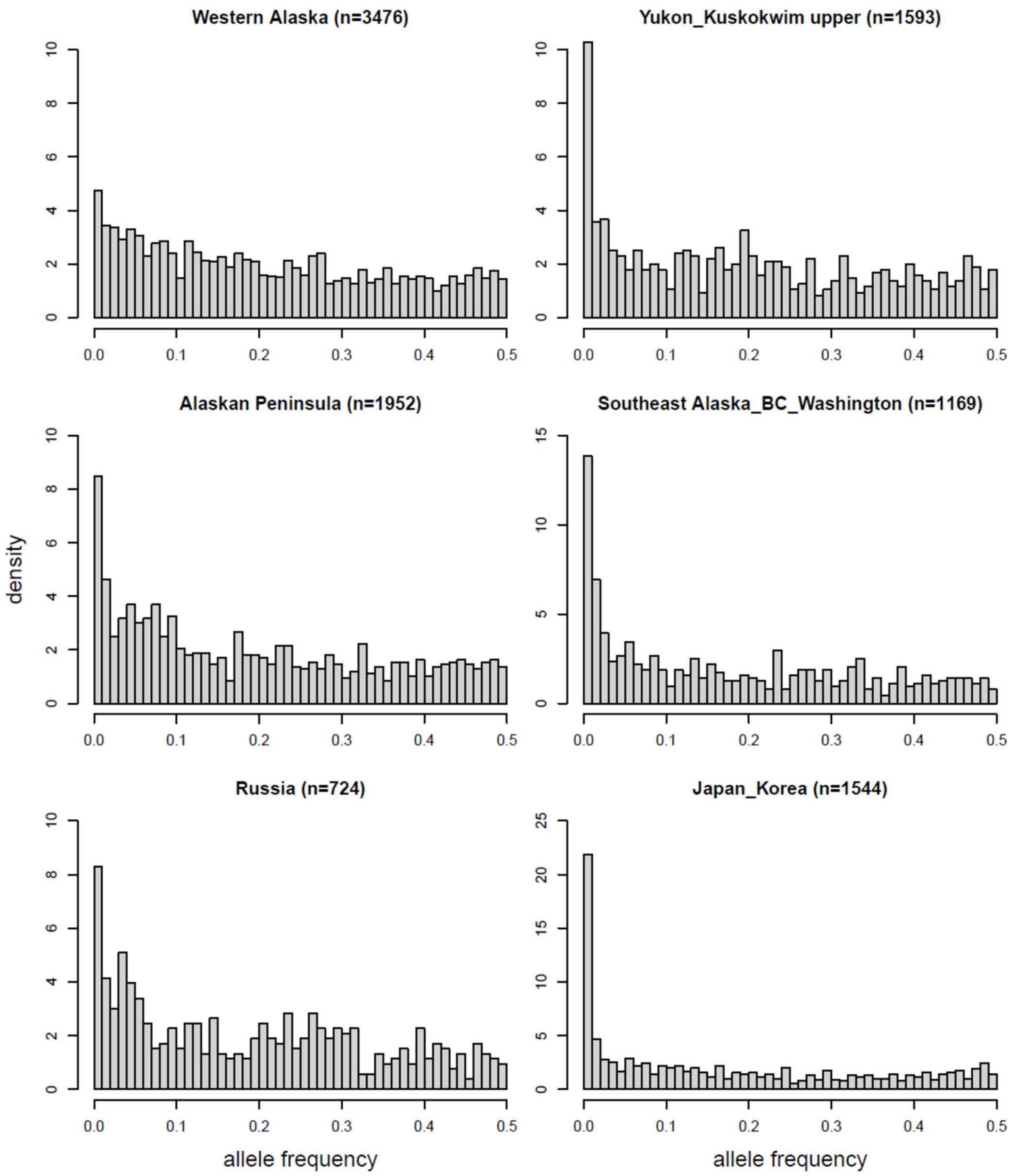


**Figure S7** Allele frequency spectra for chum salmon SNPs by six sampling areas. Data are from Seeb et al. (2011) and are given in Table S2.

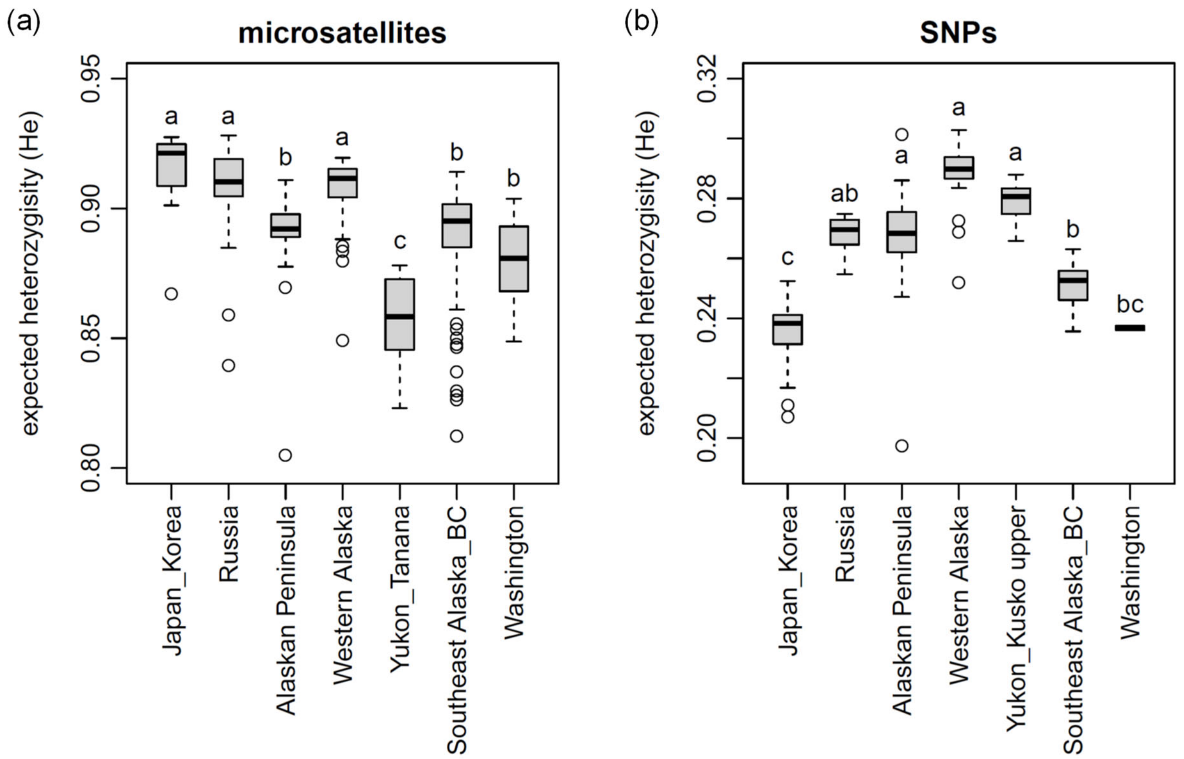


**Figure S8** Geographical distributions of expected heterozygosity (*He*) averaged over loci for (a) 10 microsatellites and (b) 53 SNPs of chum salmon. Differences between areas ($p$ <0.05) are indicated by different lowercase letters. Data are from Beacham et al. (2009a) and Seeb et al. (2011) are given in Tables S1 and S2.

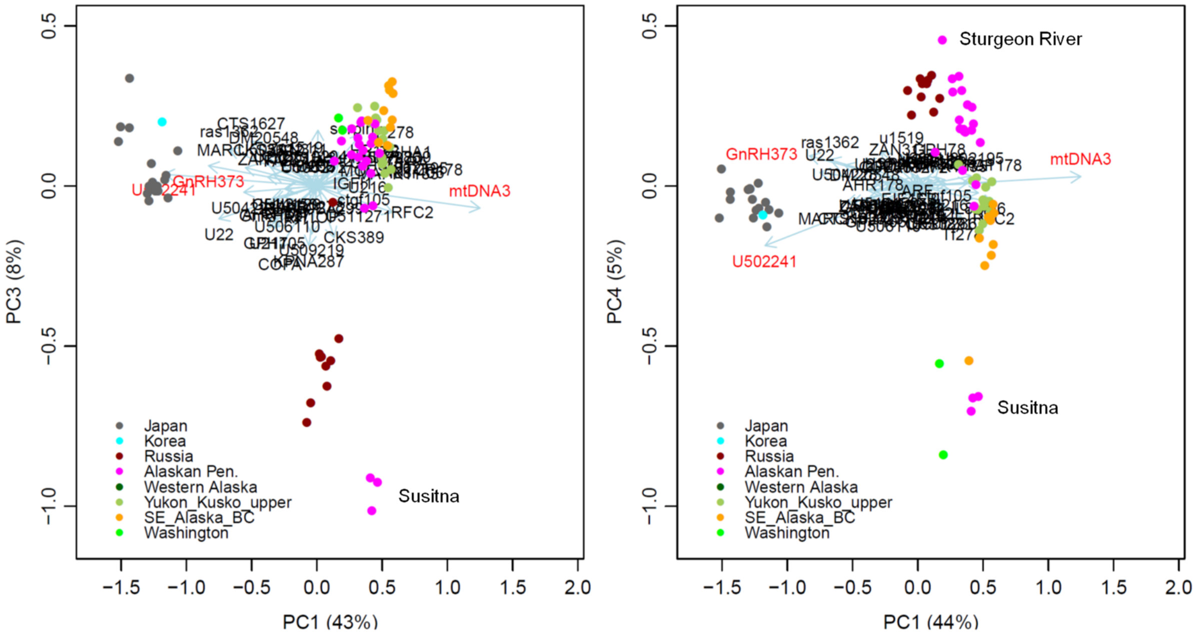


**Figure S9** Principal component analyses (PCAs) plots based on 53 SNPs and a combined mtDNA locus of chum salmon. Data are from Seeb et al. (2011) and are given in Table S2.

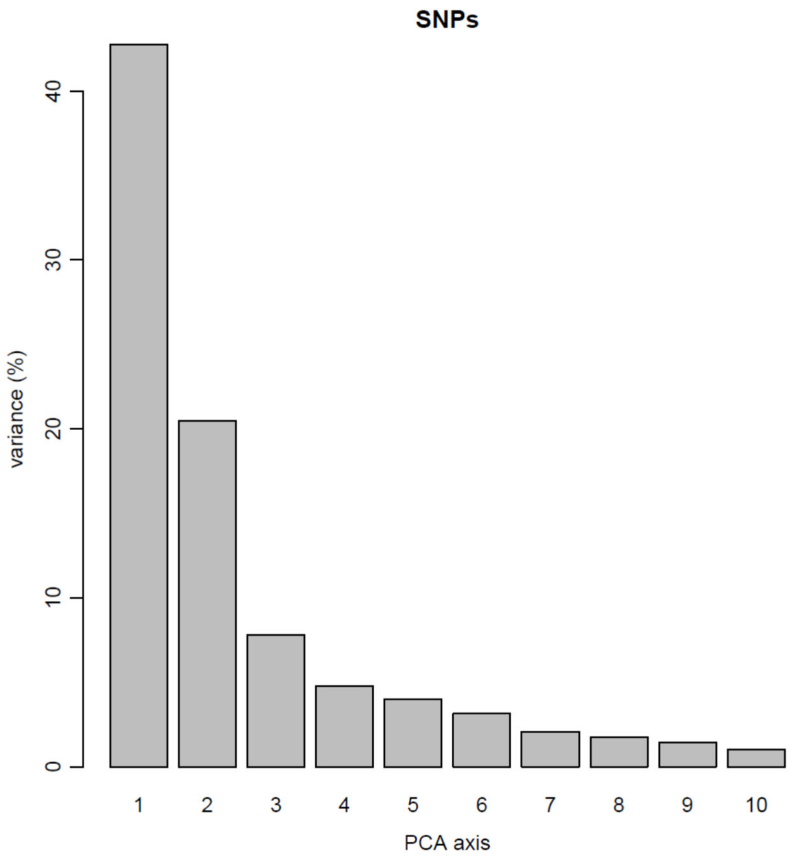


**Figure S10** Variances of principal component analyses (PCAs) based on 53 SNPs and a combined mtDNA locus of chum salmon. Data are from Seeb et al. (2011) and are given in Table S2.
